# Perturbation of alphavirus and flavivirus infectivity by components of the bacterial cell wall

**DOI:** 10.1101/2021.05.07.443110

**Authors:** Lana Langendries, Sofie Jacobs, Rana Abdelnabi, Sam Verwimp, Suzanne Kaptein, Pieter Baatsen, Lieve Van Mellaert, Leen Delang

## Abstract

The impact of the host microbiota on arbovirus infections is currently not well understood. Arboviruses are viruses transmitted through the bites of infected arthropods, predominantly mosquitoes or ticks. The first site of arbovirus inoculation is the biting site in the host skin, which is colonized by a complex microbial community that could possibly influence arbovirus infection. We demonstrated that pre-incubation of arboviruses with certain components of the bacterial cell wall, including lipopolysaccharides (LPS) of some Gram-negative bacteria and lipoteichoic acids or peptidoglycan of certain Gram-positive bacteria, significantly reduced arbovirus infectivity *in vitro*. This inhibitory effect was observed for arboviruses of different virus families, including chikungunya virus of the *Alphavirus* genus and Zika virus of the *Flavivirus* genus, showing that this is a broad phenomenon. A modest inhibitory effect was observed following incubation with a panel of heat-inactivated bacteria, including bacteria residing on the skin. No viral inhibition was observed after pre-incubation of cells with LPS. Furthermore, a virucidal effect of LPS on viral particles was noticed by electron microscopy. Therefore, the main inhibitory mechanism seems to be due to a direct effect on the virus particles. Together, these results suggest that bacteria are able to decrease the infectivity of alphaviruses and flaviviruses.

**Importance:** During the past decades the world has experienced a vast increase in epidemics of alphavirus and flavivirus infections. These viruses can cause severe diseases such as hemorrhagic fever, encephalitis and arthritis. Several alpha- and flaviviruses, such as chikungunya virus, Zika virus and dengue virus, are significant global health threats because of their high disease burden, their widespread (re-)emergence and the lack of (good) anti-arboviral strategies. Despite the clear health burden, alphavirus and flavivirus infection and disease are not fully understood. A knowledge gap in the interplay between the host and the arbovirus is the potential interaction with host skin bacteria. Therefore, we studied the effect of (skin) bacteria and bacterial cell wall components on alphavirus and flavivirus infectivity in cell culture. Our results show that certain bacterial cell wall components markedly reduced viral infectivity by directly interacting with the virus particle.

## Introduction

Arboviruses (i.e. <u>ar</u>thropod-<u>bo</u>rne viruses) are viruses transmitted through the bites of infected arthropods, predominantly mosquitoes or ticks. It is a genetically highly diverse group with more than 600 members described, of which at least 80 are known to cause disease in humans and animals (1). Due to increased travelling, climate change and adaptation of the arthropod vectors to urbanization, the geographic distribution of arboviral infections has expanded and is still spreading in many regions of the world (2). Well-known, medically important mosquito-borne viruses are chikungunya virus (CHIKV), belonging to the *Alphavirus* genus of the *Togaviridae*, and dengue virus (DENV), Zika virus (ZIKV) and yellow fever virus (YFV), belonging to the *Flavivirus* genus of the *Flaviviridae* (3). They all typically manifest first with fever and flu-like symptoms, possibly accompanied by rash or myalgia and arthralgia (4).

Although arboviruses are very diverse, they share a common characteristic: transmission via the bite of an arthropod vector into the skin. The skin is the largest organ of the human body and has a protective role against foreign organisms or toxic substances (5). The skin epidermis and dermis are colonized by a large number of microorganisms, most of which are beneficial or harmless (6). It is estimated that between 10^6^ and 10^9^ microorganisms/cm^2^ are present on the human skin (7). Microbial colonization is person-specific and depends on age, sex and environmental factors (such as clothing, use of antibiotics, soaps and cosmetics) (5, 8).

Microbiota are crucial for human health since they have an important role in the regulation of the mucosal immune system (9). Furthermore, the presence of microbiota can induce competition with other pathogens, such as pathogenic bacteria or viruses (10). Several bacterium-virus interactions have been characterized: *Lactobacillus rhamnosus* and *Bifidobacterium bifidum* bacteria have been shown to prevent rotavirus-induced diarrhea in mice by inducing inflammatory or mucosal protective factors, respectively (11, 12). Several species of the *Lactobacillus* genus inhibited replication of HIV-1 in cervico-vaginal tissue culture by producing lactic acid causing an antiviral effect (13). For influenza virus, it has been shown that lipopolysaccharides (LPS), molecules on the outer membrane of Gram-negative bacteria, activated Toll-like receptors and inhibited viral infection by direct interaction with influenza virions (14, 15). For coronaviruses (CoV) (common cold associated-CoV and MERS-CoV), it was recently described that peptidoglycan (PG) of *Bacillus subtilis* reduced coronavirus infectivity, triggered by a PG-associated cyclic lipopeptide (surfactin), which also decreased the infectivity of other enveloped viruses (e.g. ZIKV, CHIKV, Mayaro virus (MAYV), Ebola virus) (16). In contrast to the protective, antiviral effect of these microbiota, other bacteria were shown to promote viral infections. It has been described that gut bacteria enhanced viral replication, transmission and pathogenesis of enteric viruses, such as norovirus, poliovirus and reovirus. Human norovirus interacted with bacterial strains isolated from human stool, probably by binding HBGA (histo-blood group antigen) -like glycans and sialylated LPS present on bacterial cells, thereby enhancing viral entry (9, 17). Poliovirus has been shown to bind bacterial LPS, which increased virion stability and cell attachment and potentially promoted poliovirus transmission (18). Also reovirus virions directly interacted with bacteria or bacterial envelope components (LPS and PG) (19). In addition, reduced reovirus replication and pathogenesis was observed in antibiotic-treated mice (20).

Currently, little information is available on the impact of host microbiota on arbovirus infections. Oral administration of antibiotics was shown to severely aggravate flavivirus infections in mice (21). The antibiotic treatment altered the bacterial abundance in the gut and the development of optimal T cell immunity, causing detrimental effects on flavivirus-induced disease in mice (21). A comparable effect of oral antibiotic treatment on alphavirus infectivity was reported: perturbation of intestinal microbiota by the antibiotics resulted in enhanced CHIKV infection by diminishing type I interferon responses in monocytes and dendritic cells (22). These data suggest that host bacteria function as protective, anti-arboviral microorganisms. However, it is not clear yet whether host (skin) bacteria directly interact with arboviruses or whether indirect mechanisms are responsible for the antiviral effect. Therefore, we studied the effect of different bacterial cell wall components and inactivated skin bacteria on alphavirus and flavivirus infectivity *in vitro* and characterized the mechanisms of interaction.

## Material and methods

### Cells and viruses

African green monkey kidney cells (Vero cells, ATCC CCL-81) and Vero E6 cells (ATCC CRL-1586) were maintained in minimal essential medium (MEM, Gibco) supplemented with 10% fetal bovine serum (FBS, Gibco), 1% L-glutamine (Gibco), 1% sodium bicarbonate (Gibco) and 1% non-essential amino acids (NEAA, Gibco). Human skin fibroblasts (ATCC CRL-2522) were maintained in MEM supplemented with 10% FBS, 1% L-glutamine, 1% sodium bicarbonate, 1% sodium pyruvate (Gibco) and 1% NEAA. Cell cultures were maintained at 37°C in an atmosphere of 5% CO_2_ and 95%-99% relative humidity. Virus propagation and *in vitro* assays were performed using similar media but supplemented with 2% FBS (assay media).

CHIKV (Indian Ocean strain 899, passage 2 stock; GenBank FJ959103.1) is a laboratory-adapted strain (a kind gift of Prof. C. Drosten, University of Bonn, Germany). Sindbis virus (SINV, strain HRsp, GenBank J02363.1, highly passaged) and Semliki Forest virus (SFV, Vietnam strain, GenBank EU350586.1, highly passaged) belong to the historical collection of viruses at the Rega Institute of Medical Research, Belgium. MAYV (strain TC625, passage 14 stock) and ZIKV (SL1602, Suriname, passage 6 stock; GenBank KY348640.1) were obtained via the EVAg consortium (https://www.european-virus-archive.com). YFV (17D vaccine strain; Stamaril, passage 2 stock; GenBank NC_002031.1) was obtained from Sanofi Pasteur MSD, Belgium. Virus stocks were stored at -80°C. The ZIKV Suriname stock was deep sequenced. The CHIKV stock was completely sequenced by Sanger sequencing. SFV, SINV, MAYV, YFV were not sequence authenticated.

### Compounds and enzymes

The TLR4 signaling inhibitor (TAK-242; CLI-095) was purchased from Invivogen. It was dissolved at a concentration of 1 mg/ml in dimethyl sulfoxide and then 10x diluted with assay medium. For each replicate experiment, the stock concentration was freshly diluted to 5 µM. RNase A was purchased from Promega and dissolved at a concentration of 4 mg/ml in TE buffer. The RNase inhibitor (RNasin® Ribonuclease Inhibitor) was purchased from Promega and was ready to use.

### Bacteria and bacterial cell wall components

Different bacterial cell wall components, i.e. LPS of *Escherichia coli* O111:B4, *Klebsiella pneumoniae*, *Pseudomonas aeruginosa*, *Serratia marcescens,* and LTA from *Bacillus subtilis,* and PG from *Bacillus subtilis, Micrococcus luteus* and *Staphylococcus aureus*, were purchased from Sigma-Aldrich. *Klebsiella pneumoniae, Pseudomonas aeruginosa, Acinetobacter lwoffii, Corynebacterium amycolatum, Cutibacterium* (formerly *Propionibacterium*) *acnes* and *Staphylococcus aureus* belonged to the collection of the Laboratory Molecular Bacteriology (KU Leuven, Rega Institute) or were a kind gift of Prof. K. Lagrou (Laboratory Medicine, University Hospital Leuven). Bacteria were cultured overnight in 5 ml Tryptic Soy Broth (BD) or Brain Heart Infusion (BD) medium at 37°C and 180 rpm. These precultures were 1:20 to 1:50 diluted in 50 ml medium and further incubated until an OD of 0.7-1 was reached. The concentration (CFU/ml) was determined by plating the cultures on Tryptic Soy agar (BD) plates. Afterwards, bacteria were inactivated by heat or by UV radiation. For the heat-inactivation, cultures were incubated at 60°C in a shaking water bath for 20 min and were subsequently centrifuged at 1500 g for 5 min. Cell pellets were washed and finally resuspended in sterile saline. Inactivation of the bacteria was checked by plating them on Tryptic Soy agar plates. For the UV radiation, cultures were centrifuged for 5 min at 1500 g and bacterial pellets were resuspended in sterile saline to an OD600 ∼1.0. For each culture, two uncovered Petri dishes with 7.5 ml cell suspension were placed in a UVP Crosslinker CL3000 (Analytik Jena). Suspensions were 3 times irradiated with 8000 mJ/cm2 for 1 min with gentle mixing of the suspensions in between. Serial dilution and subsequent plating of the irradiated suspensions confirmed a bacterial killing effect of > 99.9% (5 to 6 log decrease in bacterial CFU count before versus after irradiation).

### Determination of viral loads after incubation with bacteria or bacterial cell wall components

Virus inoculum (10^6^ PFU/ml) was pre-incubated at 37°C for 1 or 4 h with an equal volume of one of the bacterial cell wall components (at a final concentration of 500 µg/ml for LPS of *K. pneumoniae*, *P. aeruginosa* and *S. marcescens,* and 1 mg/ml for LPS *E. coli*, LTA *B. subtilis* and PG from *B. subtilis, M. luteus and S. aureus*) or 10^10^ heat/UV-inactivated bacterial cells/ml from *K. pneumoniae, P. aeruginosa, A. lwoffii, C. amycolatum, C. acnes* and *S. aureus*. Control samples were prepared by mixing the same virus inoculum with 2% assay medium. At the end of the incubation time, the infectious virus progeny in the virus/bacterial cell wall component, virus/bacteria mixtures or the control were quantified by end-point titration on Vero cells.

Briefly, cells were seeded in 96-well microtiter plates (at a density of 2.5×10^4^ cells/well for Vero cells, 2×10^4^ cells/well for Vero E6 cells and 1.2×10^4^ cells/well for CRL-2522 cells) and were allowed to adhere overnight. The next day, 3 parallel 10-fold serial dilutions of the virus/bacterial component or virus/bacteria mixtures were prepared in the plates. After the specified incubation time for each virus (3 days in case of alphaviruses, 7 days for flaviviruses), the cells were examined microscopically for virus-induced cytopathogenic effect (CPE). A well was scored positive if any traces of virus-induced CPE were observed compared to the uninfected controls. The TCID_50_/ml was calculated using the method of Reed and Muench (23) and is defined as the virus dose that would infect 50% of the cell cultures. Limits of quantification (LOQs) were determined as the lowest viral loads that could be quantified using this method.

For the experiment with the TLR4-inhibitor, virus/LPS mixtures, which were pre-incubated at 37°C for 1 h, were added to the cells (Vero or skin fibroblasts), where after the TLR4-inhibitor TAK-242 was added at an end concentration of 5 µM.

To study the effect on intra- and extracellular viral RNA levels, Vero cells were pre-seeded at 2.5×10^4^ cells/well in a 96-well plate (BD Falcon). The next day, after 4 h of pre-incubation, cells were infected in triplicate with either a virus/bacterial cell wall component mixture (as described above) or control mixture. At 2 h post-infection (pi), the inoculum was removed and cells were washed three times with assay medium. At 24 h pi, extracellular viral RNA was extracted using the NucleoSpin RNA virus kit (Macherey-Nagel) and intracellular viral RNA using the E.Z.N.A Total RNA Kit I (Omega Bio-tek), according to the manufacturer’s protocols. Viral RNA levels were quantified by quantitative reverse transcription PCR (qRT-PCR).

### Immunofluorescence assay

Skin fibroblasts (2.4x10^4^ cells/well) were seeded in 8-well chamber slides (Ibidi) that were pre-treated with poly-L-lysin (Merck) to improve cell attachment. The next day, cells were pre-incubated with TLR4 inhibitor (TAK-242; 5 µM) for 1 h and exposed to a mixture of LPS of *K. pneumoniae* (end concentration of 50 µg/ml) and TAK-242 (end concentration of 5 µM) for 2 h. Controls without LPS or without TAK-242 were processed alongside. After exposure, cells were fixed with 3.7% formaldehyde (Sigma-Aldrich) for 20 min. The day after, cells were permeabilized with 0.5% Triton X-100 (Sigma-Aldrich) for 10 min and were blocked with 3% bovine serum albumin (BSA; Sigma) for 30 minutes, where after cells were stained with NF-κB p65 antibody (1:200 dilution in 1% BSA in PBS; Santa Cruz Biotechnology) in the dark for 1 h. Goat anti-mouse IgG Alexa Fluor 488 (1:500 dilution in 1% BSA in PBS; Invitrogen) was used as the secondary antibody. Nuclei were stained with Hoechst (1:1000 dilution in PBS; Invitrogen) in the dark for 20 min. Cellular localization of NF-κB was determined with a Leica TCS SP5 laser scanning confocal microscope (Leica Microsystems), using a HCX PL APO 63.0x (NA:1.20) water immersion lens.

### Virucidal assay

Virus (10^6^ PFU/ml) was pre-incubated at 37°C with LPS of *K. pneumoniae* or *P. aeruginosa* (at a final concentration of 500 µg/ml) or 2% assay medium for 1 h, after which 1 μ mg/ml, Promega) was added to the reaction mixture. As a positive control, extracted viral RNA was treated with RNase A. After 1 h incubation at 37°C, 1 µL of RNAse inhibitor (40 U/µl, Promega) was added to stop the reaction. The viral RNA in each sample was extracted using the NucleoSpin RNA virus kit (Macherey-Nagel) and quantified by qRT-PCR.

### Quantitative reverse transcription PCR (qRT-PCR)

The sequences of primers used in qRT-PCR were for CHIKV: forward primer: 5′☐CCGACTCAACCATCCTGGAT☐3′, reverse primer: 5′☐GGCAGACGCAGTGGTACTTCCT☐3′(24); for SFV: forward primer: 5′-GCAAGAGGCAAACGAACAGA-3′, reverse primer: 5′-GGGAAAAGATGAGCAAACCA-3′ (25); for ZIKV: forward primer: 5’-CCGCTGCCCAACACAAG-3’, reverse primer: 5’-CCACTAACGTTCTTTTGCAGACAT-3’ (26). The probe sequences were 5’-FAM-TCCGACATCATCCTCCTTGCTGGC-TAMRA-3’ (CHIKV) (24) and 5’-FAM-AGCCTACCT-ZEN-TGACAAGCAATCAGACACTCAA-IABkFQ-3’ (ZIKV) (26). One-step, quantitative RT-PCR was performed for CHIKV and ZIKV in a total volume of 25 µl, containing 13.94 µl RNase free water (Promega), 6.25 µl master mix (Eurogentec), 0.375 µl of forward primer, 0.375 µl of reverse primer [final concentration of each primer: 150 nM (CHIKV); 900 nM (ZIKV)], 1 µl of probe [final concentration: 400 nM (CHIKV); 200 nM (ZIKV)], 0.0625 µl reverse transcriptase (Eurogentec) and 3 µl RNA sample. For SFV, the reaction mixture (20 µl) contained: 4.75 µl RNAse free water, 10 µl SYBR green master mix (Bio-Rad), 1 µl of forward primer, 1 µl of reverse primer (final concentration of each primer 500 nM), 0.25 µl reverse transcriptase (Eurogentec) and 3 µl sample. The qRT-PCR reaction was performed with the Applied Biosystems 7500 Fast Real-Time PCR System using the following conditions for CHIKV and ZIKV: 30 min at 48°C, 10 min at 95°C, followed by 40 cycles of 15 s at 95°C and 1 min at 60°C. For SFV, the following conditions were used: 10 min at 50°C, 3 min at 95°C, followed by 40 cycles of 15 s at 95°C and 30 s at 60°C.

For quantification, standard curves were generated each run using 10-fold dilutions of cDNA of CHIKV nsP1 and SFV nsP3 or viral RNA isolated from the ZIKV Suriname virus stock (virucidal assay) or a synthesized gene block containing ZIKV E protein (assay to determine effect of LPS on viral RNA levels). Limits of detection (LODs) were determined as the lowest viral loads that could be detected by the qRT-PCR assay in 95% of experiments (27).

### Transmission electron microscopy (TEM)

SFV (6x10^8^ PFU/ml) was pre-incubated at 37°C with LPS of *K. pneumoniae* or *P. aeruginosa* (final concentration of 500 µg/ml) or LPS of *E. coli* (final concentration of 1 mg/ml) for 1 h. Pre-incubation of SFV with LPS of *S. marcescens* (final concentration of 500 µg/ml) was performed for different incubation times (0, 5, 10, 15, 20, 25, 30 min or 1 h). SFV was pre-incubated at 37°C with heat-inactivated *K*. *pneumoniae* or *S. aureus* bacteria (10^10^ cells/ml) for 1 h. Viruses were inactivated by glutaraldehyde (Electron Microscopy Sciences; final concentration 1.25%) at room temperature for 30 min. Formvar-carbon coated 400-mesh copper grids (Ted Pella) were first glow-discharged to improve adsorption efficiency, and were then placed on 20 µl of sample for 5 min, after which excess of sample was removed by blotting on filter paper. In order to increase virus density on the grid, grids were further incubated 4 times for 1 minute with intermittent blotting. Subsequently, grids were washed by contact with 2 drops of milliQ water and were negatively stained with 20 µl of 1% uranyl acetate (Electron Microscopy Sciences) for 1 min, after which excess was removed using filter paper. After drying, the grids were examined using a transmission electron microscope (JEOL JEM1400), operated at 80 kV and equipped with an EMSIS Quemesa 11 Mpxl camera. The number of viruses that could be detected at a magnification of 4000 x was counted for each incubation time at 6 randomly picked positions on the grid (each position represented a surface of 132 µm^2^ of the grid).

### *Ex vivo* skin culturing

Mouse skin was collected under the approval of the Ethical Committee of the University of Leuven (licenses P071/2019, M020/2020) following institutional guidelines approved by the Federation of European Laboratory Animal Science Associations (FELASA). In-house-bred / female, > 20 weeks old AG129 mice (deficient in both IFN-α/β and IFN-γ receptors) were sacrificed by intraperitoneal injection of Doléthal (Vétoquinol). The back of the mice was shaven and dorsal skin was collected and immediately covered in Dulbecco’s Modified Eagle Medium (DMEM; Gibco) supplemented with 10% FBS, 1% penicillin/streptomycin (Gibco) and 0.1% gentamycin (Sigma) on ice. Next, subcutaneous adipose tissue was removed and the dorsal skin was cut into pieces of approximately 1 cm². After a wash step in PBS, skin samples were individually placed (epidermis-side up) in a 12-well (BD Falcon) plate and incubated in supplemented DMEM at 37°C until infection.

### Infection of skin explants with virus pre-incubated with LPS

SFV (1x10^4^ PFU/ml) was pre-incubated with an equal volume of LPS of *K. pneumoniae*, *P. aeruginosa* and *S. marcescens* (at a final concentration of 500 µg/ml) at 37°C for 4 h. Control samples were prepared by adding an equal volume of 2% assay medium to the SFV inoculum. The SFV-LPS mixtures were added to the skin explants and after 2 h of incubation at 37°C, the inoculum was removed. Skin explants were washed 3 times in PBS and were again incubated in supplemented DMEM. At day 2 pi, the skin tissue was transferred to a Precellys tube (Bertin Instruments) containing 2% assay medium. Skin tissues were homogenized in two cycles at 7600☐rpm with a 20☐s interval, using an automated homogenizer (Precellys24, Bertin Instruments). After centrifugation (15 000 rpm, 15 min, 4°C), supernatant was collected and infectious virus was quantified by end-point titrations.

### Statistics

All data were analysed using Graphpad Prism 8.3.1. The results of the virucidal assays were statistically analysed using a nonparametric Two-way ANOVA with Sidak’s correction for multiple comparisons. The effect of LPS on viral RNA levels was analysed using the Mann Whitney U test (nonparametric t-test). All other results were analysed by Kruskal-Wallis tests (nonparametric One-way ANOVA). Statistical significance threshold was assessed at *p* values of <0.05. Statistical details are described in the figure legends.

## Results

### CHIKV infection is reduced by certain components of the bacterial cell wall

To study the effect of bacterial cell wall components on arbovirus infectivity, different cell wall structures (LPS of *K. pneumoniae*, and LTA or PG of *B. subtilis*) were incubated with CHIKV and viral infectivity was determined by end-point titration on Vero cells. LPS of K. *pneumoniae* completely blocked viral infectivity, whereas a modest inhibitory effect of ∼1.5 log_10_ reduction was observed with PG of *B. subtilis* (Fig. 1A). LTA of *B. subtilis* resulted in less than 1 log_10_ reduction in infectious virus levels. The complete disruption of CHIKV infectivity by LPS of *K. pneumoniae* was also confirmed at viral RNA level, both intracellularly and in the supernatant (Fig. 1B). As the structures of LPS can significantly differ between different bacteria and as the preparation method of PG might affect the composition, we explored whether the observed reduction in virus infectivity by LPS and PG was shared by LPS/PG from other bacterial species. To this end, the effect of a panel (Table 1) of LPS from Gram-negative bacteria (*P. aeruginosa, E. coli, S. marcescens)* and PG of Gram-positive bacteria (*S. aureus, M. luteus)* on CHIKV infectivity was evaluated (Fig. 1C). Clear differences in the ability to decrease CHIKV infection in Vero cells were observed between LPS and PG of different bacterial species: incubation with most LPS and PG resulted only in a modest inhibitory effect (1-2 log_10_ reduction), whereas no effect was observed with PG of *S. aureus* and *M. luteus* (< 1 log_10_ reduction). These data suggest that the reduction of CHIKV infectivity is not widely shared by LPS and PG molecules of all bacteria.

**Figure 1.**
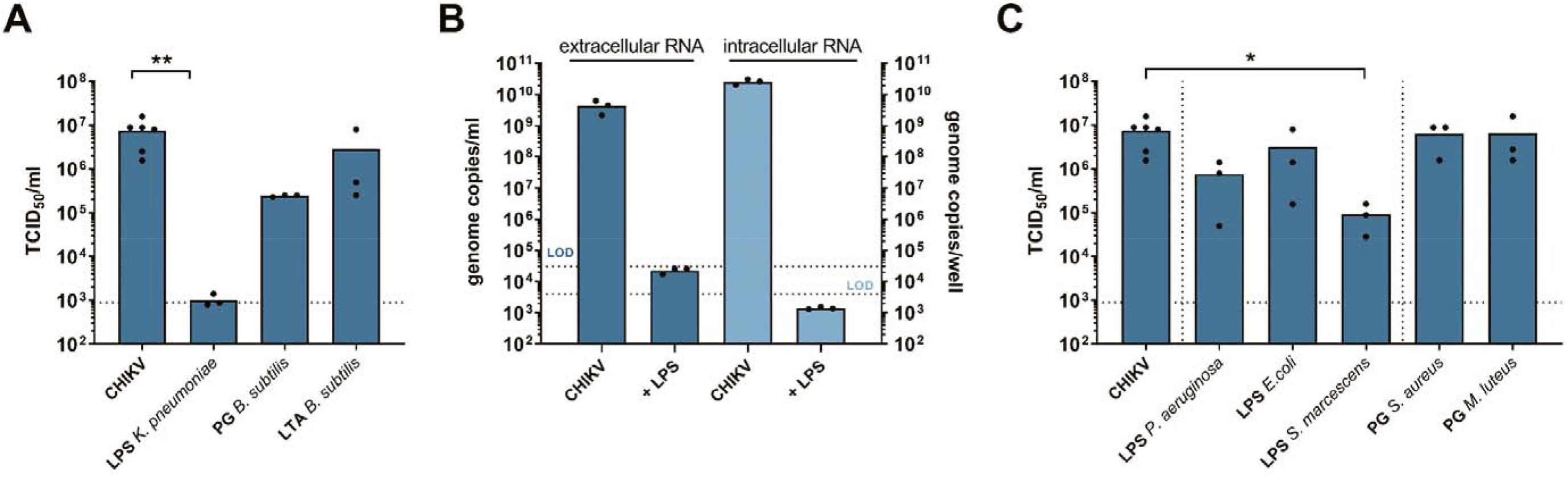
CHIKV infectivity is reduced by different components of the bacterial cell wall. **(A)** Infectivity of CHIKV determined by end-point titrations on Vero cells (TCID_50_/ml). CHIKV was incubated with different bacterial cell wall components (LPS of *K. pneumoniae*, PG or LTA of *B. subtilis*) at 37°C for 4 h. **(B)** CHIKV RNA levels, intracellular (genome copies/well) and extracellular (genome copies/ml supernatant), determined by qRT-PCR. After 4 h incubation at 37°C of virus with LPS of *K. pneumoniae*, mixtures were added to Vero cells and viral RNA levels were determined at 24 h pi. Individual data points are shown with the height of the bar representing the mean value. Statistical significance was assessed with a Mann Whitney U test. The dotted lines represent the LODs. **(C)** Infectivity of CHIKV after incubation with an expended panel of LPS (of *P. aeruginosa*, *E. coli* and *S. marcescens*) and PG (of *S. aureus* and *M. luteus*) at 37°C for 4 h. **(A,C)** Individual data points are shown with the height of the bar representing the mean value. Statistical significance was assessed with a Kruskal-Wallis test. Significantly different values are indicated by asterisks: *p* < 0.05: *, *p* <0.005: **. The dotted line represents the limit of quantification (LOQ). TCID, tissue culture infectious dose; LPS, lipopolysaccharide; PG, peptidoglycan; LTA; lipoteichoic acid.

**Table 1.** Bacterial species used in this study.

| GRAM STAINING | PHYLUM | BACTERIAL SPECIES | SKIN COLONIZATION |
| --- | --- | --- | --- |
| **Negative** | *Proteobacteria* | *Acinetobacter lwoffii* | resident of the skin flora (45–47) |
|  |  | *Escherichia coli* | rare pathogenic skin colonizer (48) |
|  |  | *Klebsiella pneumoniae* | rare resident of the skin flora (49)<br>+ pathogenic skin colonizer (50) |
|  |  | *Pseudomonas aeruginosa* | resident of the skin flora (51) +<br>pathogenic skin colonizer (52, 53) |
|  |  | *Serratia marcescens* | rare pathogenic skin colonizer (54) |
| **Positive** | *Actinobacteria* | *Corynebacterium amycolatum* | resident of the skin flora +<br>pathogenic skin colonizer (55, 56) |
|  |  | *Micrococcus luteus* | resident of the skin flora (57–59) |
|  |  | *Cutibacterium acnes* | resident of the skin flora +<br>pathogenic skin colonizer (60) |
|  | *Firmicutes* | *Bacillus subtilis* | rare resident of the skin flora (61) |
|  |  | *Staphylococcus aureus* | resident of the skin flora +<br>pathogenic skin colonizer (62, 63) |

### Certain LPS and PG reduce the infectivity of different alphaviruses and flaviviruses

To find out whether this inhibitory effect is a broad-spectrum phenomenon amongst arboviruses, the panel of bacterial cell wall components was evaluated against arboviruses of two virus families: SFV, SINV and MAYV of the *Alphavirus* genus (*Togaviridae*) and ZIKV and YFV of the *Flavivirus* genus (*Flaviviridae*). Comparable results were observed for most alphaviruses (Fig. 2A) and flaviviruses (Fig. 2B), except for PG of *B. subtilis* and *S. aureus*, which were able to completely inhibit ZIKV infection (Fig. 2B) and LPS of *S. marcescens* which inhibited SFV, SINV (Fig. 2A) and ZIKV replication (Fig. 2B) in Vero cells. Furthermore, LPS of *K. pneumoniae* could not completely reduce MAYV (Fig. 2A) and ZIKV (Fig. 2B) infectivity, whereas LPS of *P. aeruginosa* inhibited SFV and SINV infectivity to the limit of detection in Vero cells (∼4 and ∼3.5 log_10_ reduction in TCID_50_/ml, respectively) (Fig. 2A).

**Figure 2.**
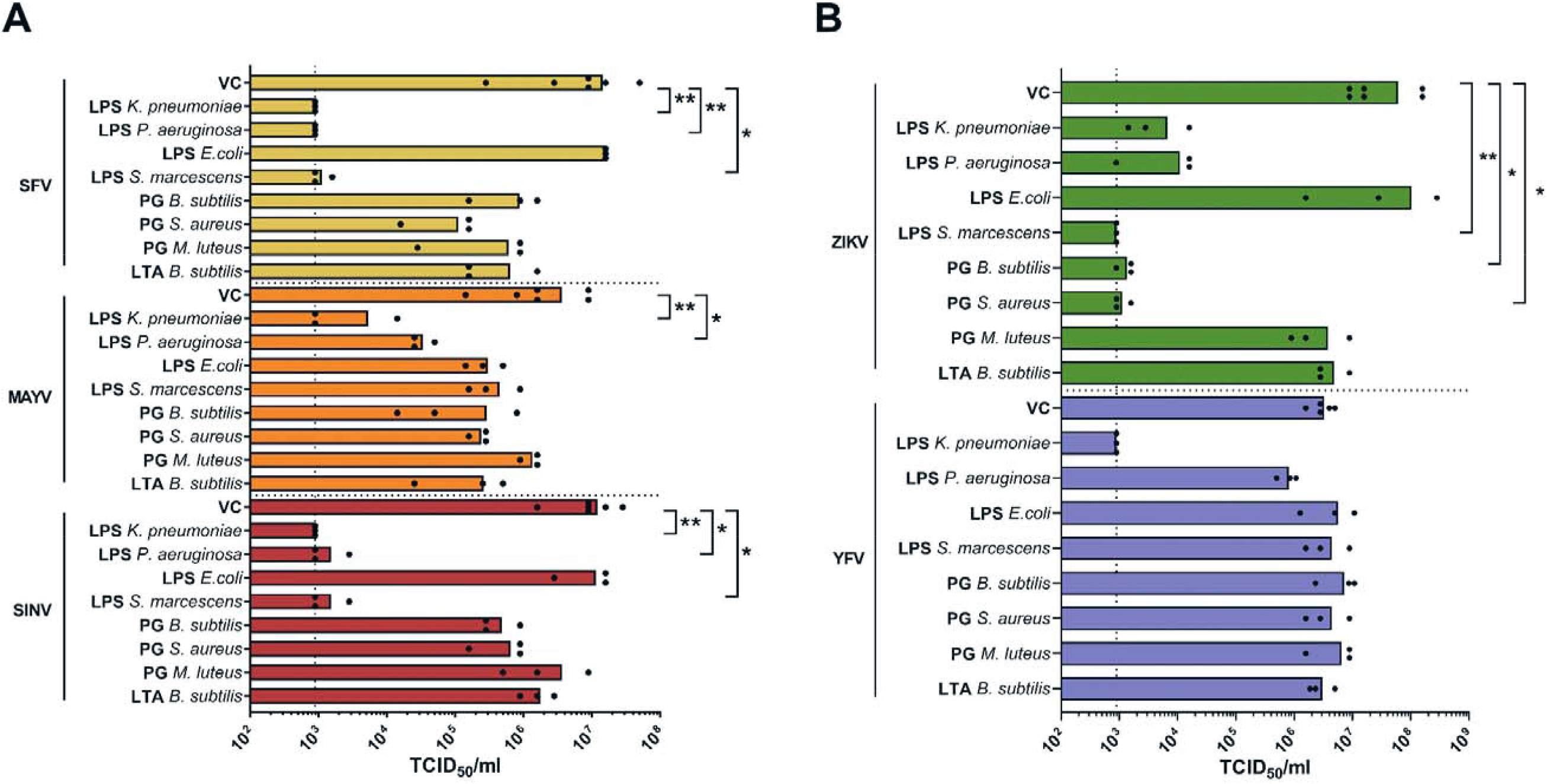
Components of the bacterial cell wall decrease the infectivity of several alphaviruses and flaviviruses. (A-B) Infectivity of **(A)** alphaviruses (SFV, MAYV and SINV) and **(B)** flaviviruses (ZIKV and YFV) quantified by end-point titrations on Vero cells (TCID_50_/ml). Viruses were incubated with a panel of LPS, PG and LTA of different bacterial species at 37°C for 4 h. Individual data points are shown with the height of the bar representing the mean value. Statistical significance was assessed with a Kruskal-Wallis test. Significantly different values are indicated by asterisks: *p* < 0.05: *, *p* <0.005: **. The dotted line represents the LOQ. TCID, tissue culture infectious dose; VC, virus control; LPS, lipopolysaccharide; PG, peptidoglycan; LTA; lipoteichoic acid.

### Specific LPS and PG reduce alphavirus infectivity in human skin fibroblasts and *ex vivo* mouse skin

To investigate whether virus inhibition would also be observed in other cell types, we assessed CHIKV infectivity after incubation with LPS or PG in human skin fibroblasts. LPS of *K. pneumoniae* completely blocked CHIKV infectivity in skin fibroblasts (Fig. 3A), similar to what was observed in Vero cells. For LPS and PG of other bacterial species, there were minor variations in the reductions in infectivity between Vero cells and skin fibroblasts. However, LPS of *S. marcescens* completely blocked CHIKV infectivity in skin fibroblasts, in contrast to the effect observed in Vero cells (∼2 log_10_ decrease).

**Figure 3.**
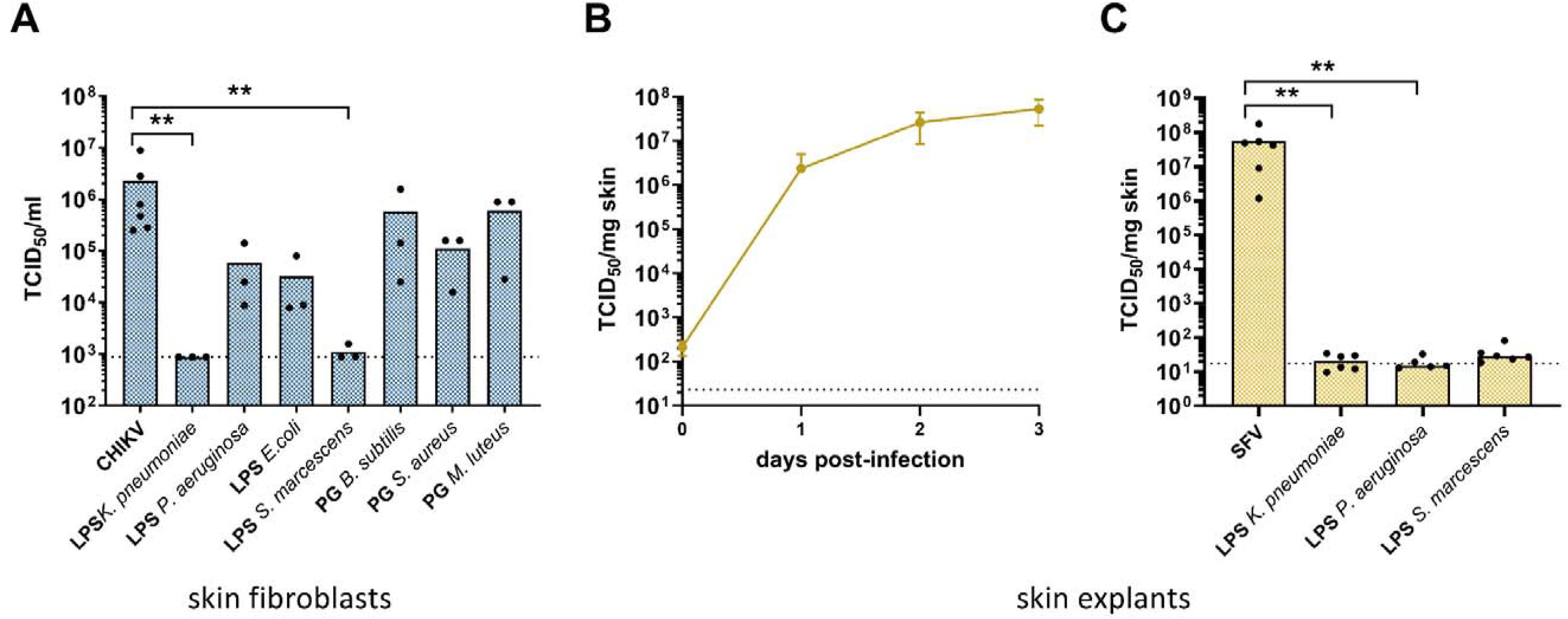
Specific LPS and PG reduce alphavirus infectivity in human skin fibroblasts and *ex vivo* mouse skin. **(A)** Infectivity of CHIKV assessed by end-point titrations on skin fibroblasts (TCID_50_/ml). CHIKV was incubated with a panel of LPS, PG and LTA at 37°C for 4 h. **(B)** Replication kinetics of SFV in *ex vivo* cultured mouse skin tissue (1x10^4^ PFU/ml). Infectious progeny virus was quantified at different time points (1, 2 and 3 d pi) by end-point titrations on Vero cells. Data shown are mean values ± SD from four individual data points. **(C)** Infectivity of SFV in mouse skin explants. SFV was incubated with LPS of *K. pneumoniae*, *P. aeruginosa* or *S. marcescens* at 37°C for 4 h and was added to skin explants. At day 2 pi, infectious virus was quantified by end-point titrations on Vero cells. **(A, C)** Individual data points are shown with the height of the bar representing the mean value. Statistical significance was assessed with a Kruskal-Wallis test. Significantly different values are indicated by asterisks: p <0.005: **. The dotted line represents the LOQ. TCID; tissue culture infectious dose; LPS, lipopolysaccharide; PG, peptidoglycan

Next, we determined the effect of LPS on viral infectivity in *ex vivo* cultured mouse skin. First, the replication kinetics of SFV were defined in the skin explants (Fig. 3B), showing that high viral titers could be reached at day 2 and day 3 pi. Next, we incubated SFV with different LPS and determined the infectious virus titer in the skin. Similar to what was observed *in vitro* (Fig. 2), LPS of *K. pneumoniae*, *P. aeruginosa* and *S. marcescens* inhibited SFV infectivity to the limit of detection (Fig. 3C).

### Alphavirus and flavivirus infectivity is reduced by specific bacteria

We next evaluated whether complete bacteria could also affect the infectivity of arboviruses. To this end, we selected a panel of gram-negative (*K. pneumoniae, P. aeruginosa* and *A. lwoffii*) and gram-positive bacteria (*S. aureus, C. amycolatum, C. acnes*) (Table 1). The Gram-positive *Staphylococcus*, *Corynebacterium* and *Cutibacterium* spp. are among the most abundant colonizers of the skin (5). The most reported Gram-negative skin bacteria are the *Acinetobacter* spp. (28, 29). After inactivation of the bacteria by heating, viruses (CHIKV, SFV and ZIKV) were incubated with these bacterial species. A modest inhibitory effect was observed with *K. pneumoniae*, *P. aeruginosa* and *A. lwoffii* on CHIKV, and with *K. pneumoniae, P. aeruginosa, A. lwoffii, C. amycolatum* and *C. acnes* on SFV (Fig. 4A). For ZIKV on the other hand, the inhibition by *K. pneumoniae*, *P. aeruginosa* and *A. lwoffii* was more pronounced (∼2-2.5 log_10_ reduction in TCID_50_/ml). To investigate whether the heat-inactivation procedure could be the reason for the differences between the effects by LPS and the bacteria, LPS of *K. pneumoniae* was subjected to the same heating procedure. Following incubation with heated LPS, CHIKV infectivity was still completely reduced (Fig. 4B), confirming that LPS is a heat-stable cell wall component (15, 30). Additionally, we evaluated the effect of three bacterial species that were inactivated by UV radiation. A similar inhibitory trend was observed, compared to bacteria that were heat-inactivated (Fig. 4C): *P. aeruginosa* resulted in the most pronounced inhibition, followed by *K. pneumoniae*, whereas *S. aureus* had the smallest effect on CHIKV infectivity.

**Figure 4.**
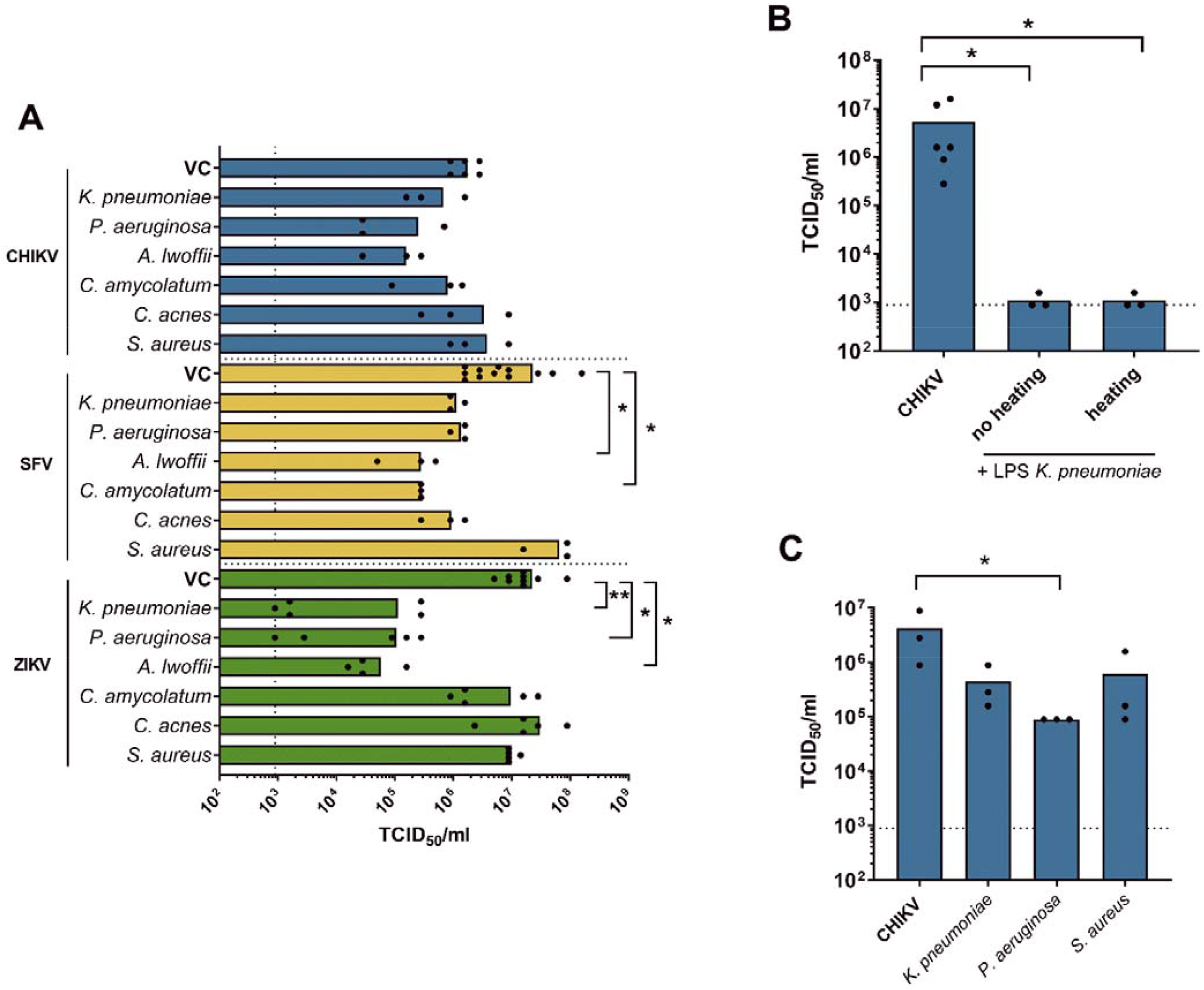
Alphavirus and flavivirus infectivity is reduced by certain inactivated bacteria. **(A)** CHIKV, SFV and ZIKV infectivity after incubation with Gram-negative (*K. pneumoniae*, *P. aeruginosa, A. lwoffii*) or Gram-positive (*C. amycolatum*, *C. acnes*, *S. aureus*) heat-inactivated bacteria at 37°C for 4 h. Viral titers were determined by end-point titrations on Vero cells (TCID_50_/ml). **(B)** CHIKV infectivity after incubation at 37°C for 1 h with LPS of *K. pneumoniae* or with LPS of *K. pneumoniae* that was pre-heated at 60°C for 20 min. **(C)** CHIKV infectivity after incubation with UV-inactivated *K. pneumoniae*, *P. aeruginosa,* and *S. aureus* bacteria at 37°C for 4 h. Viral titers were determined by end-point titrations on Vero cells (TCID_50_/ml). **(A-C)** Individual data points are shown with the height of the bar representing the mean value. Statistical significance was assessed with a Kruskal-Wallis test. Significantly different values are indicated by asterisks: *p* < 0.05: *, *p* <0.005: **. The dotted line represents the LOQ. TCID; tissue culture infectious dose; VC, virus control; LPS, lipopolysaccharide.

### Disruption of virus infectivity is time- and dose-dependent

To determine how fast the inhibition of infection occurs, CHIKV, SFV and ZIKV were incubated with LPS of *K. pneumoniae* at 37°C for 2 min, 10 min, 30 min, 1 h or 4 h (Fig. 5A). LPS of *K. pneumoniae* was selected for this purpose, as this molecule was the most potent of the evaluated panel of bacterial cell wall components. For both alphaviruses, a short incubation time with LPS was sufficient to achieve complete inhibition (CHIKV: 30 min; SFV: 10 min). For ZIKV on the other hand, a fast drop in infectivity was initially observed (∼3 log_10_), but at least 60 min of incubation was required to result in a maximal inhibition of ∼3.5 log_10_ in TCID_50_/ml.

**Figure 5.**
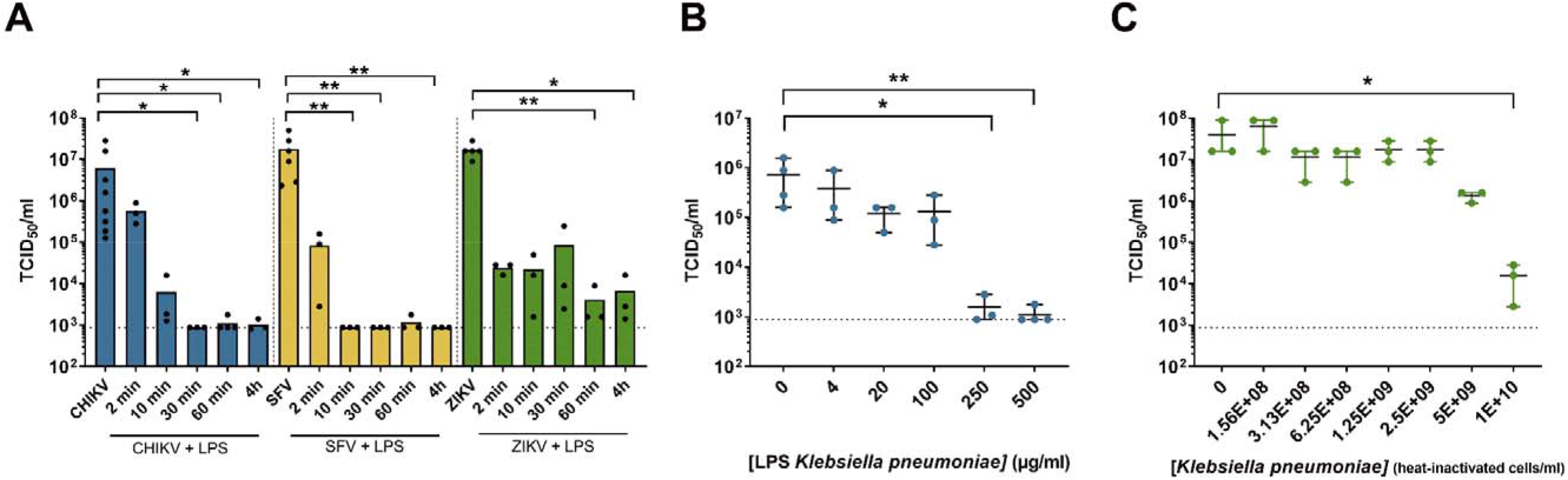
Disruption of virus infectivity is time- and dose-dependent. **(A)** Time-dependent effect of LPS of *K. pneumoniae*: infectivity of CHIKV, SFV and ZIKV was determined by end-point titrations on Vero cells (TCID_50_/ml), after incubation at 37°C with LPS of *K. pneumoniae* for different time periods (2 min, 10 min, 30 min, 1 h, 4 h). Individual data points are shown with the height of the bar representing the mean value. **(B)** Dose-dependent effect of LPS of *K. pneumoniae*: CHIKV infectivity after incubation with serial dilutions of LPS of *K. pneumoniae* (0 µg/ml, 4 µg/ml, 20 µg/ml, 100 µg/ml, 250 µg/ml, 500 µg/ml) at 37°C for 1 h. **(C)** Dose-dependent effect of heat-inactivated *K. pneumoniae* bacteria: ZIKV infectivity after incubation with serial dilutions of *K. pneumoniae* (0, 1.56E+08, 3.13E+08, 6.25E+08, 1.25E+09, 2.5E+09, 5E+09, 1E+10 heat-inactivated cell/ml) at 37°C for 4 h. **(B, C)** Individual data points and the mean value are shown. **(A-C)** Statistical significance was assessed with a Kruskal-Wallis test. Significantly different values are indicated by asterisks: *p* < 0.05: *, *p* <0.005: **. The dotted line represents the LOQ. TCID; tissue culture infectious dose; LPS, lipopolysaccharide.

To determine the minimum inhibitory concentration of LPS of *K. pneumoniae* and heat-inactivated *K. pneumoniae* bacteria, serial dilutions were incubated with CHIKV and ZIKV, respectively, and infectivity was determined by end-point titration on Vero cells. The effective concentration of LPS of *K. pneumoniae* that inhibited 90% of virus-induced cell death (EC_90_) was 167 µg/ml and a concentration of 250 µg/ml was necessary to completely block the virus (Fig. 5B). A concentration of 1x10^10^ heat-inactivated bacterial cells/ml of *K. pneumoniae* resulted in a ∼3 log_10_ reduction in ZIKV infectivity (Fig. 5C). The EC_90_ value of *K. pneumonia*e bacteria was 4.17x10^9^ cells/ml. In contrast to LPS, complete inhibition with heat-inactivated bacteria was not achieved.

### Is the reduced virus infectivity cell- or virus-dependent?

The reduction of virus infectivity might be due to a direct interaction between LPS and the virus or by an effect of LPS on cellular immune pathways (or a combination of both). Comparable effects in skin fibroblasts and Vero cells were observed (Fig. 1 and 3A), except for LPS of *S. marcescens*, which caused a more prominent inhibition of CHIKV infection in skin fibroblasts. Therefore, we pre-incubated LPS of *S. marcescens* and *K. pneumoniae* with cells (Vero cells or skin fibroblasts) or virus (CHIKV) and determined viral infectivity by end-point titration (Fig. 6A). LPS of *K. pneumoniae* was enclosed as well, since this was the only LPS that could inhibit all tested viruses in this study. For both LPS, inhibition of CHIKV infectivity was more pronounced when LPS was pre-incubated with virus, both in Vero cells and in skin fibroblasts, suggesting that the main inhibitory mechanism is due to a direct interaction between LPS and the viral particle.

**Figure 6.**
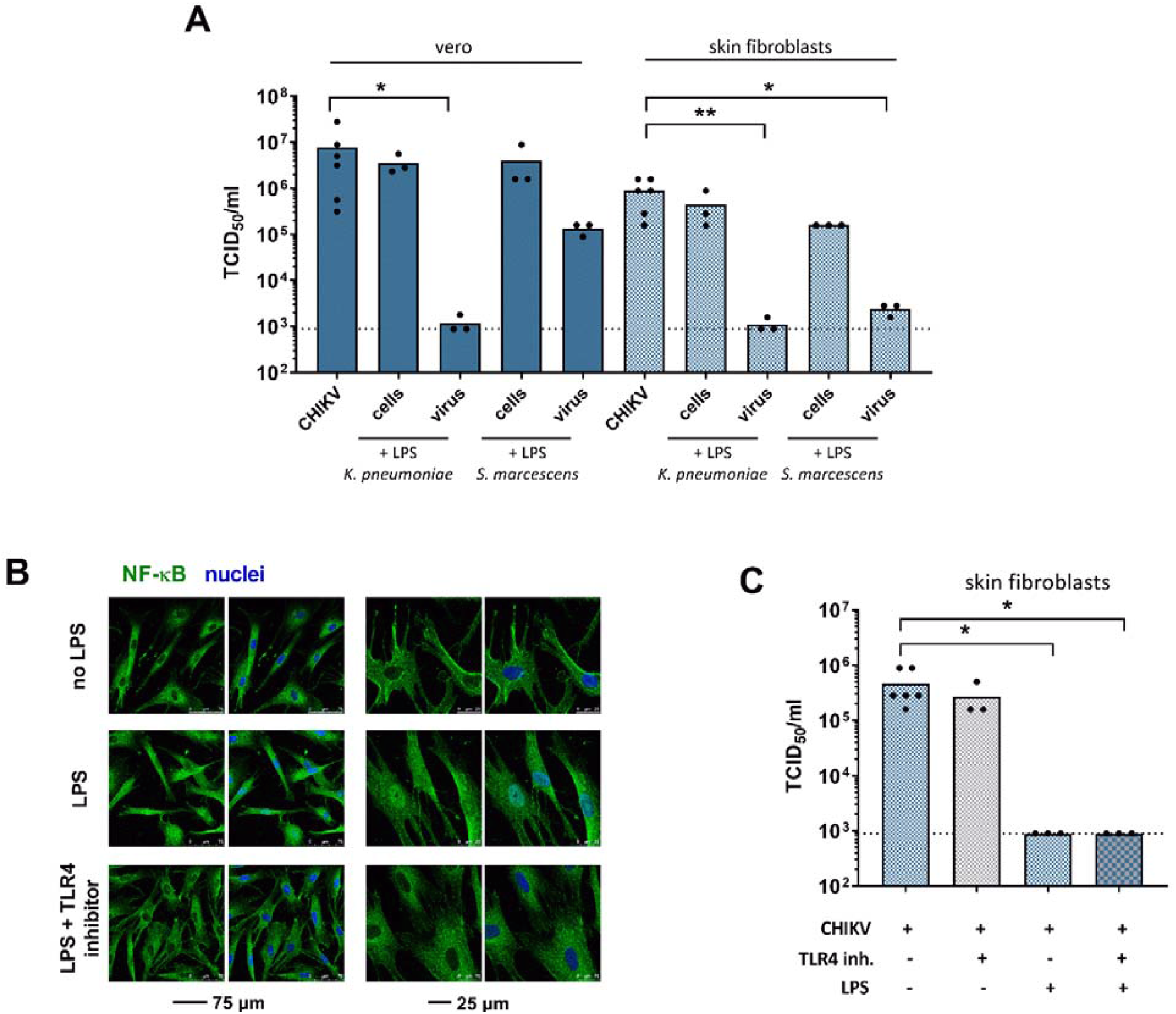
The reduction in virus infectivity is probably not due to a cell-dependent effect. **(A)** CHIKV infectivity after pre-incubation of cells (Vero cells or skin fibroblasts) with LPS of *K. pneumoniae*/*S. marcescens* or after pre-incubation of virus with LPS of *K. pneumoniae/S. marcescens* at 37°C for 1 h. Viral loads were determined both on Vero cells and skin fibroblasts by end-point titrations. **(B)** NF-κB p65 (green) was visualized in skin fibroblast cells cultured for 2 h in the absence (panel 1) or presence of LPS of *K. pneumoniae* (panel 2) or LPS of *K. pneumoniae* and a TLR4 inhibitor (panel 3) using confocal fluorescence microscopy. Nuclei were stained with Hoechst (blue). **(C)** CHIKV infectivity determined by end-point titrations on skin fibroblasts (TCID_50_/ml) following incubation for 1 h with TAK-242, a TLR4 inhibitor and/or LPS of *K. pneumoniae*. **(A, C)** Individual data points are shown with the height of the bar representing the mean value. Statistical significance was assessed with a Kruskal-Wallis test. Significantly different values are indicated by asterisks: *p* <0.05: *, *p* <0.005: **. The dotted line represents the LOQ. TCID, tissue culture infectious dose; LPS, lipopolysaccharide.

It has been described that bacteria can activate an LPS-dependent Toll-like receptor 4 (TLR4) immune pathway, resulting in an antiviral effect (14). TLR4 is expressed in human skin fibroblasts (31) and LPS can induce the activation of the TLR4-NF-κB immune pathway in mouse fibroblasts (32). To investigate whether the observed viral inhibition in skin fibroblasts might be due to the stimulation of this immune pathway by LPS, we first confirmed the activation of TLR4 by LPS in human skin fibroblasts by immunofluorescence staining of NF-κB. Upon incubation with LPS, the NF-κB protein was translocated from the cytoplasm to the nucleus (Fig.6B panel 1-2), proving that the skin fibroblasts indeed express TLR4. The TLR4 inhibitor TAK-242 was able to block the activation of the TLR4-NF-κB pathway in the skin fibroblasts, since the fluorescent signal was only visible in the cytoplasm (Fig 6B panel 3). Finally, we assessed viral infectivity upon incubation with LPS in the presence of TAK-242. TAK-242 alone had no adverse effect on CHIKV infection in the skin fibroblasts (Fig. 6C). When added to cell cultures infected with LPS pre-incubated virus, TAK-242 did not affect the inhibitory effect of LPS on CHIKV infection (Fig. 6C), suggesting that the effect on viral infectivity is not due to TLR4 activation by LPS.

### LPS exerts a virucidal effect on alphaviruses and flaviviruses

To determine how virus infectivity is reduced by incubation with LPS, virucidal assays were performed with two alphaviruses (CHIKV and SFV) and one flavivirus (ZIKV). Incubation of CHIKV virus particles with LPS of *K. pneumoniae* in addition of RNase resulted in a decrease of ∼4 log_10_ in viral genome copies, whereas incubation with LPS of *P. aeruginosa* did not (Fig. 7A). This was in line with our previous experiments measuring viral infectivity (Fig. 1B). On the other hand, SFV RNA levels were reduced with ∼2.5 log_10_ after incubation with LPS of both bacteria and RNase (Fig. 7B). For ZIKV, we noticed a more modest effect: viral RNA levels were reduced to a lesser extent after incubation with LPS, compared to the positive RNA control (Fig. 7C).

**Figure 7.**
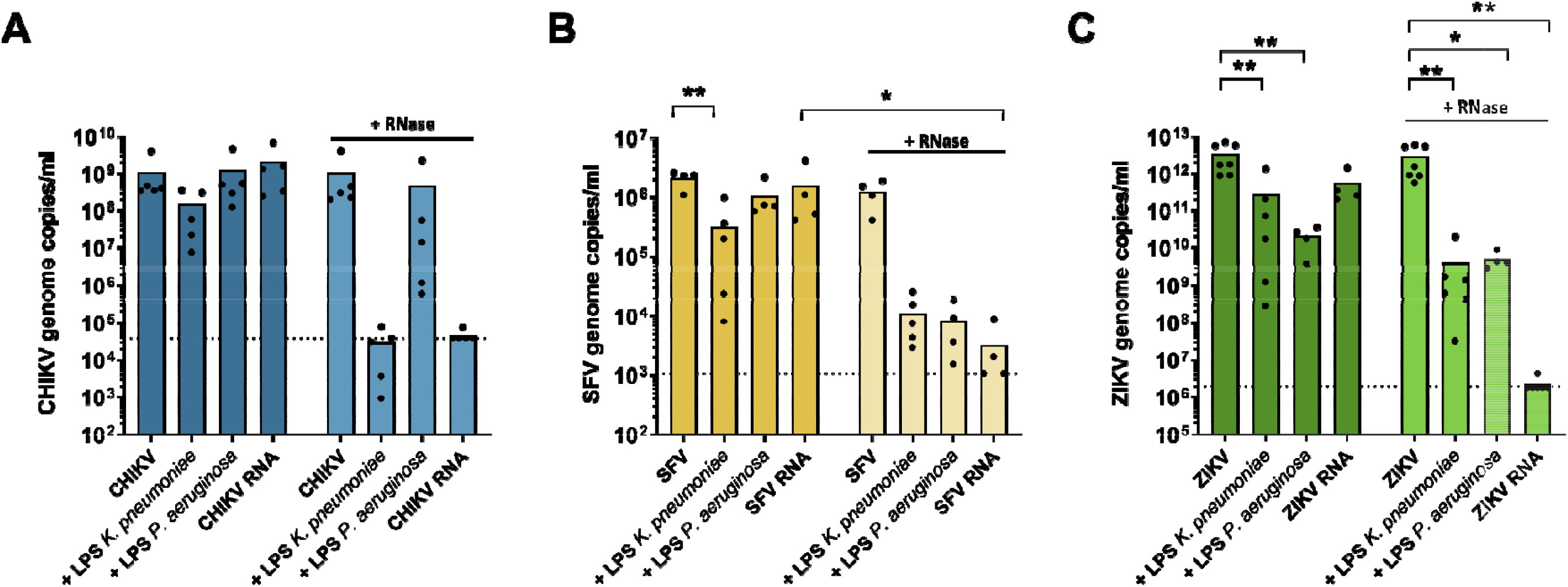
LPS exerts a virucidal effect on alphaviruses and flaviviruses. **(A)** CHIKV, **(B)** SFV or **(C)** ZIKV genome copies/ml determined by qRT-PCR. After 1 h incubation at 37°C of virus with LPS of *K. pneumoniae* or *P. aeruginosa*, RNase was added and viral RNA was quantified. Individual data points are shown with the height of the bar representing the mean value. Statistical significance was assessed with a nonparametric Two-way ANOVA with Sidak’s correction for multiple comparisons. Significantly different values are indicated by asterisks: *p* <0.05: *, *p* <0.005:**. The dotted lines represent the limits of detection (LOD). LPS, lipopolysaccharide.

To confirm that the virucidal effect was caused by a direct interaction between LPS and the virus, we imaged SFV particles in the absence and presence of LPS by TEM. LPS of *K. pneumoniae*, *P. aeruginosa* and *S. marcescens* were selected as these LPS completely inhibited SFV infectivity in cell culture (Fig. 2A). The structures of LPS of *K. pneumoniae* and *P. aeruginosa* were difficult to distinguish from virus particles as the LPS consisted of multiple globular shapes (Fig. 8A-D). LPS of *S. marcescens* on the other hand had a filamentous structure, which was easier to distinguish from virus particles (Fig. 9A). LPS of *E. coli* was included as a negative control (Fig. 9C-D). After incubation with LPS of *E. coli* for 1 h, the structure of the SFV particles was similar to the structure of untreated viruses (Fig. 10A). In contrast, upon incubation with LPS of *S. marcescens*, the morphology of the virus particles changed with longer incubation times (Fig. 10A). In addition, the number of viruses detected at a magnification of 4000 x decreased with longer incubation times (Fig. 10B). Following 30 minutes of incubation, no or very few viruses could be detected anymore (Fig 9B).

**Figure 8.**
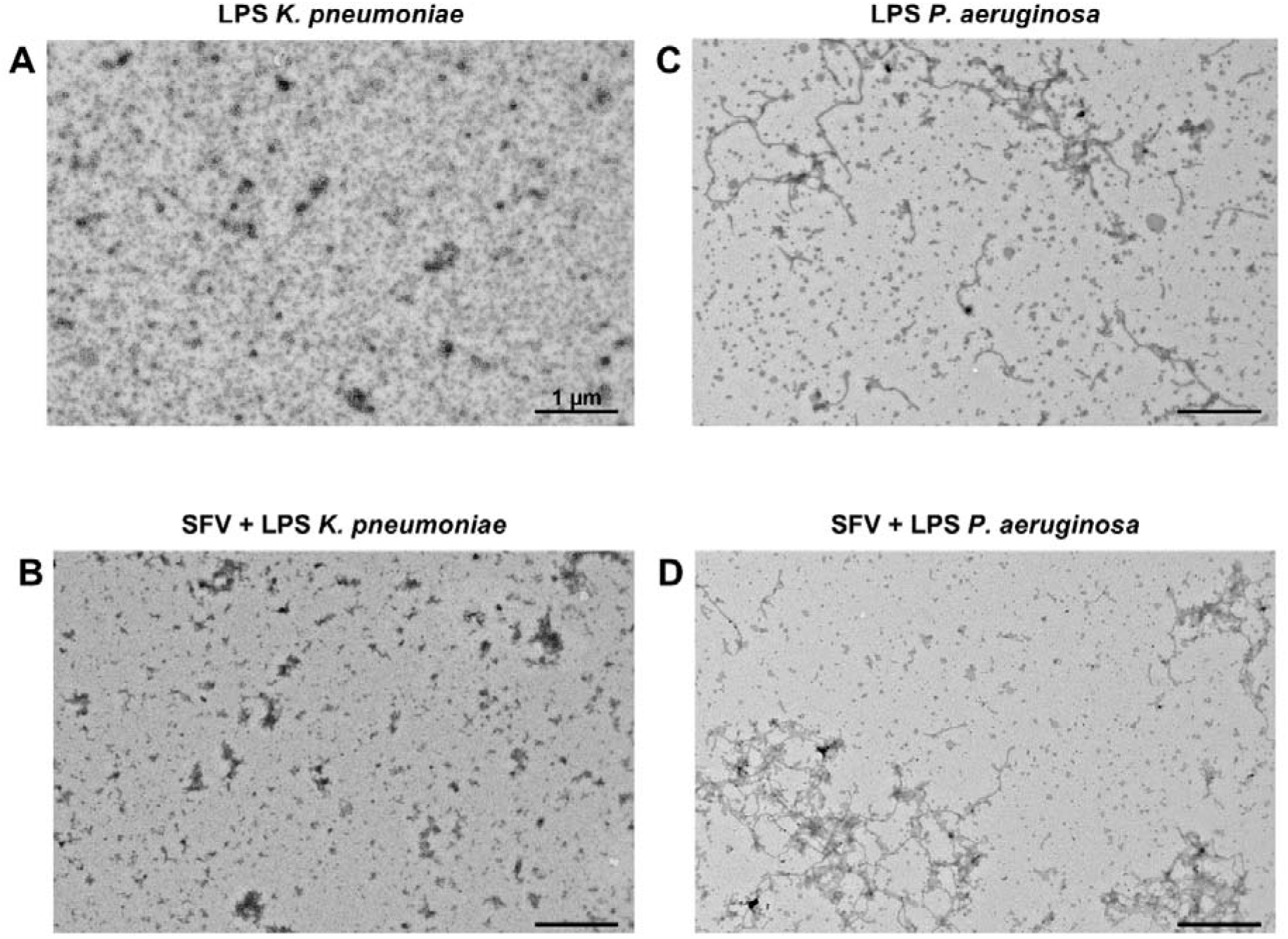
TEM images of LPS of K. pneumoniae and P. aeruginosa. **(A)** TEM image of LPS of *K. pneumoniae* at a magnification of 4000 x. **(B)** TEM image of LPS of *K. pneumoniae* incubated with SFV at 37°C for 1 h. The image was taken at a magnification of 4000 x**. (C)** TEM image of LPS of *P. aeruginosa* at a magnification of 4000 x. **(D)** TEM image of LPS of *P. aeruginosa* incubated with SFV at 37°C for 1 h. The image was taken at a magnification of 4000 x.

**Figure 9.**
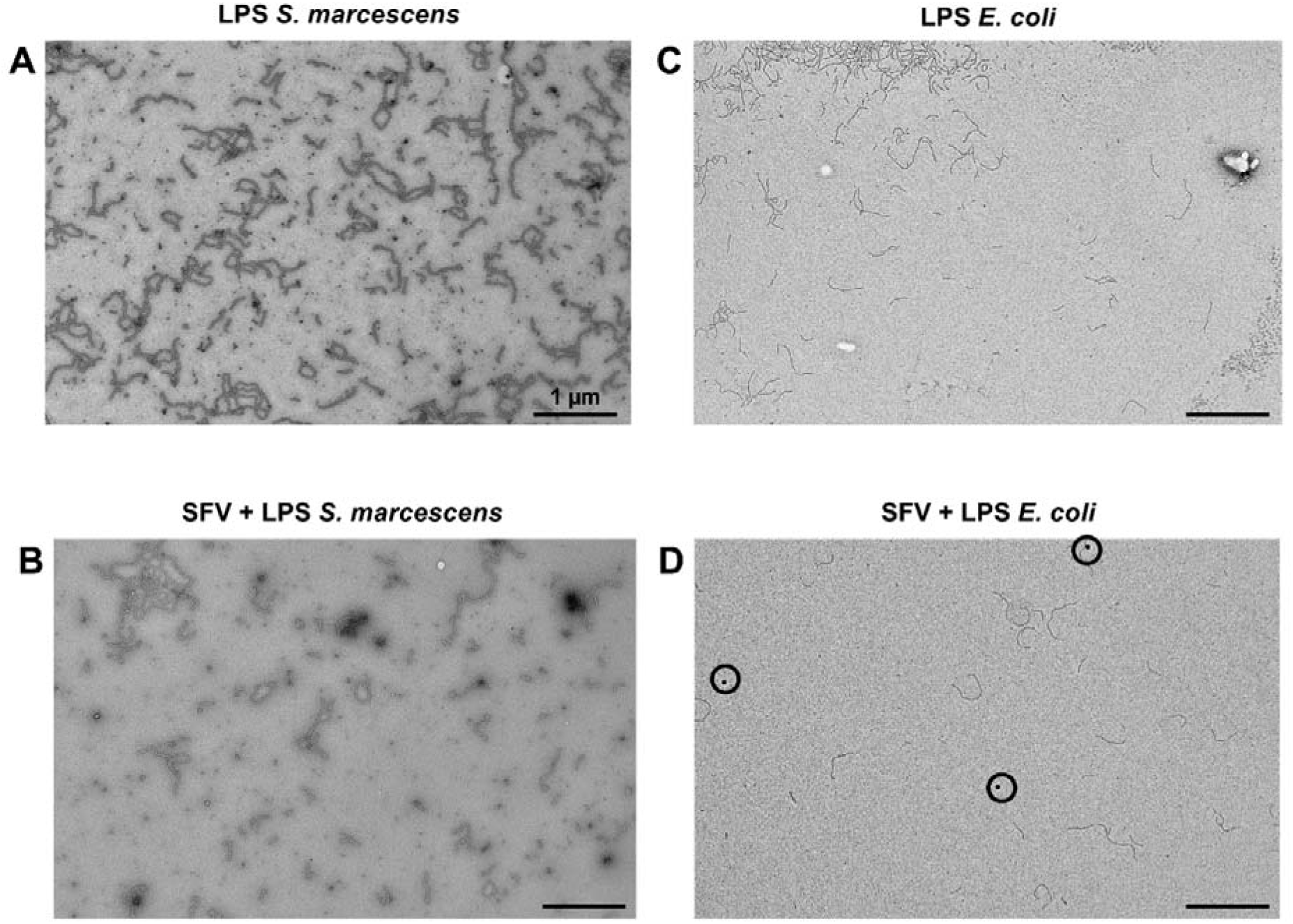
TEM images of LPS of S. marcescens and E. coli. **(A)** TEM image of LPS of *S. marcescens* at a magnification of 4000 x. **(B)** TEM image of LPS of *S. marcescens* incubated with SFV at 37°C for 1 h. The image was taken at a magnification of 4000 x. **(C)** TEM image of LPS of *E. coli* at a magnification of 4000 x. **(D)** TEM image of LPS of *E. coli* incubated with SFV at 37°C for 1 h. The image was taken at a magnification of 4000 x. SFV particles are indicated by black circles.

**Figure 10.**
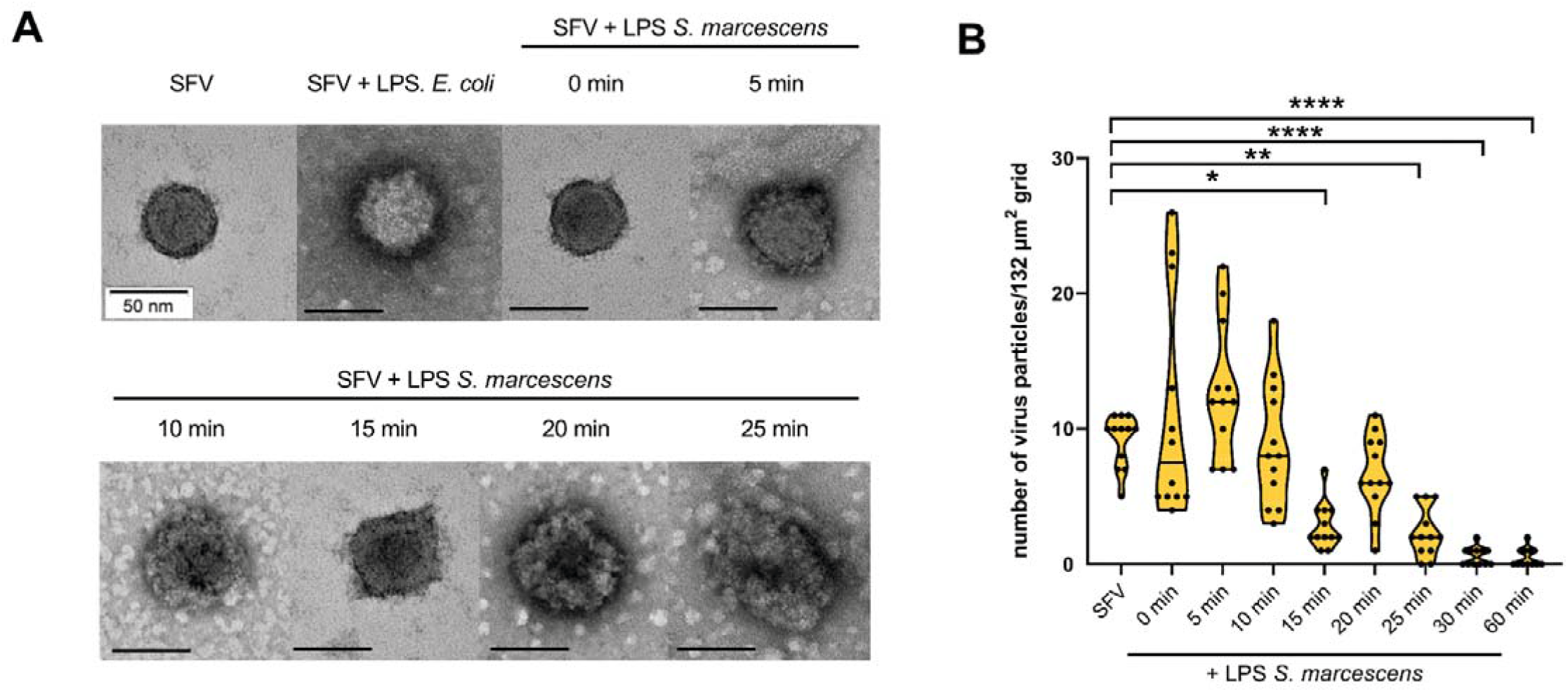
Virucidal effect of LPS of *S. marcescens* on SFV visualized by TEM. **(A)** TEM images of SFV or SFV incubated with LPS of *E. coli* at 37°C for 60 min or SFV incubated with LPS of *S. marcescens* at 37°C for 0, 5, 10, 15, 20 and 25 min. Images were taken at a magnification of 100 000 x. The black bars represent a scale of 50 nm. LPS, lipopolysaccharide. **(B)** Number of SFV particles for different incubation times with LPS of *S. marcescens*. Twelve images were captured for each condition at 4000 x at random positions on the grids (representing 132 µm^2^) and the number of virions was counted. Data are from 2 independent experiments and are presented in violin plots showing individual data points and medians. Statistical analysis was performed using the Kruskal-Wallis test. Significantly different values are indicated by asterisks: *p* <0.05: *, *p* <0.005:**, *p* <0.0001:****.

Next, we determined whether a similar effect was observed with heat-inactivated bacteria. After 1 h of incubation with *K. pneumoniae*, virus particles were absent in the TEM images (Fig. 11A), whereas virus particles were still present after incubation with *S. aureus* (Fig 11B). The TEM data thus strengthened our hypothesis that specific LPS or bacteria exert a virucidal effect on alphaviruses and flaviviruses.

**Figure 11.**
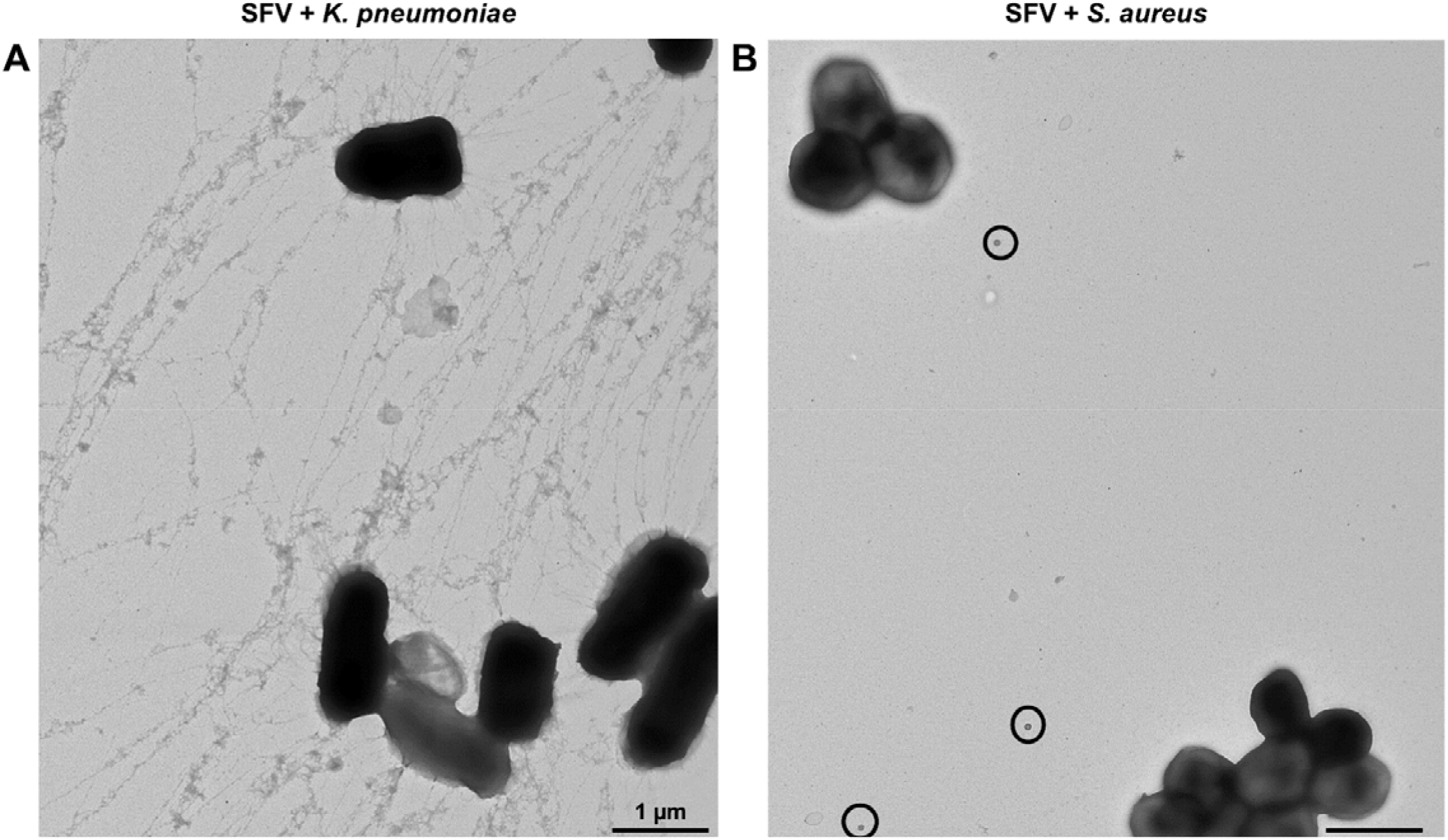
TEM images of SFV after incubation with *K. pneumoniae* and *S. aureus*. **(A)** TEM image of *K. pneumoniae* bacteria incubated with SFV at 37°C for 1 h. The image was taken at a magnification of 4000 x. **(B)** TEM image of *S. aureus* bacteria incubated with SFV at 37°C for 1 h. The image was taken at a magnification of 4000 x. SFV particles are indicated by black circles.

## Discussion

Complex interplays between viruses and host microbiota have recently gained more interest. Especially viruses that colonize the gastrointestinal tract have been studied the most due to the vast microbial community at the site of the initial virus infection (33). Currently, little research is performed regarding the role of host microbiota in arbovirus infectivity. Since arboviruses are first inoculated in the skin during the bite of an arthropod, the purpose of this research was to characterize interactions and involved mechanisms between arboviruses and the microbiota that colonize the skin. First, we tested the effect of different bacterial cell wall components (LPS, PG and LTA) on alphavirus and flavivirus infectivity. Our data indicated that not all tested bacterial cell wall structures were equally potent in inhibiting virus infectivity. Especially between the tested LPS, major differences were detected, ranging from no antiviral activity (e.g. LPS *E. coli*) to a complete inhibitory effect (e.g. LPS of *K. pneumoniae*). The structural variability of LPS is generally attributed to the polysaccharide part, particularly to the O-antigen (34). Furthermore, it has also been described that subtle chemical variations in the lipid A structure of LPS can cause drastic changes in LPS activity (35), potentially resulting in differential potencies to inhibit viral replication. Moreover, there were differences in potency of the same LPS/PG against different viruses. The general trend was the inhibitory effect of LPS of *K. pneumoniae* (on all tested viruses), LPS of *P. aeruginosa* (except on CHIKV) and LPS of *S. marcescens* (on SFV, SINV and ZIKV). All these bacteria are Gram-negative *Proteobacteria*, while the human skin (dermis and epidermis) is believed to be quantitatively dominated by the Gram-positive *Cutibacteria*, *Corynebacteria* (both *Actinobacteria*), and *Staphylococcus* species (*Firmicutes*) (28, 29, 36). Nevertheless, approximately 23% of the bacteria found on the skin are Gram-negative *Proteobacteria* or *Bacteriodetes* (37), whereby *P. aeruginosa* (a *Proteobacterium* used in our panel) was detected in human skin samples of the forearm (28). Furthermore, a more diverse population of bacteria resides in dry skin regions (5, 8), with a greater prevalence of Gram-negative β-*Proteobacteria* and *Flavobacteriales* (37).

Next, we studied the effect of complete bacteria, as this better mimics the real-life situation. Arboviruses are inoculated by the mosquito into the dermal layer of the skin (1). It has been shown that bacteria are not only present on the outermost epidermal layer, but also in the dermis (6, 29), making it possible that arboviruses and bacteria encounter in the dermal skin. *Pseudomonas spp*., *C. acnes*, *C. amycolatum*, *A. lwoffii* and *Staphylococcus spp.* have previously been identified in the dermis (6, 29). Therefore, we included several of these bacteria in this study. The effect of complete bacteria was however less pronounced, compared to the effect of LPS: only for ZIKV was there a clear inhibitory effect of *A. lwoffii*, *K. pneumoniae* and *P. aeruginosa* on infectivity. The Gram-positive skin bacteria resulted in a modest viral inhibition, although the magnitude of inhibition was similar to the inhibition previously observed for influenza virus (15) by some bacterial isolates. In contrast, the Gram-negative skin bacterium *A. lwoffi* resulted in the highest inhibitory effect, which suggests that mainly Gram-negative bacteria reduce viral infectivity. Similar effects were observed with bacterial cell wall components (Fig. 2): LPS of Gram-negative *K. pneumoniae*, *P. aeruginosa* and *S. marcescens* completely inhibited viral infectivity, whereas most of the PG of Gram-positive bacteria did not exert a clear inhibitory effect. A possible reason for the more modest inhibition seen with the complete, heat-inactivated bacteria could be that the concentrations of purified LPS used in our assay, were much higher than the concentrations LPS present in complete bacteria. For our assays, we used a concentration of 500 µg/ml LPS, based on the dose-dependent effect of LPS. To the best of our knowledge, it has not been determined yet which concentrations of LPS are representative for the human skin.

LPS concentrations can widely differ between organs: for example for the gut, an estimated concentration of 1 g of LPS has been reported, resulting in mg/ml concentrations, whereas only 10-20 pg/ml has been detected in plasma of healthy volunteers (38). Therefore, further studies concerning the concentrations of LPS in the human skin are necessary to determine the physiological relevance of our research. Furthermore, it is possible that the purified, commercially available LPS can have other effects than the natural cell wall-associated form in the complete, heat-inactivated bacteria, where the lipid A part is embedded in the membrane, hidden from the surface. Another reason could be that the used heat-inactivation procedure may have altered the LPS molecules or the structure of the outer membrane that encloses these molecules. However, LPS has been shown to be heat-stable (15, 30) and this was confirmed in our experiment by the fact that the inhibitory effect of LPS was not lost after the heating procedure. Furthermore, UV-inactivated bacteria caused the same inhibitory trend as seen with the heat-inactivated bacteria, confirming that the inactivation procedure did not significantly influence the results.

We next determined whether the reduced virus infectivity by LPS was due to a cell-dependent effect. Skin fibroblasts were included, as they are the first cells that are infected by alphaviruses after the arthropod bite (39) and it is very likely that they are also infected in the early stages of flavivirus infections (40, 41). Viral infectivity was only significantly reduced when LPS was pre-incubated with the virus and not when LPS was pre-incubated with the cells, suggesting that the viral inhibition is not due to a cell-dependent effect. A well-characterized cellular interaction of LPS is the activation of Toll-like receptor 4, resulting in increased innate immunity and triggering inflammation and adaptive immune responses (35). To exclude that the viral inhibition was caused by binding of LPS to TLR4, which subsequently activates the TLR4 immune pathway, a TLR4 antagonist was added. In the presence of the TLR4 inhibitor, a similar reduction in viral infectivity was observed by LPS, indicating that the inhibitory effect was not due to the binding and activation of TLR4. Together with other data of this study (Fig. 6), these results suggested that LPS directly interacted with alphaviruses and flaviviruses. We showed that disruption of viral infectivity was at least partially due to a virucidal effect: there was a decrease in viral RNA when LPS was incubated with the virus particles. This effect was clearly smaller with the flavivirus ZIKV, compared to alphaviruses CHIKV and SFV. To further confirm the hypothesis of a direct effect between alphaviruses and bacteria, SFV particles were imaged by TEM. Our data suggested that disruption of the virus occured upon incubation of SFV with LPS of *S. marcescens* and heat-inactivated *K. pneumoniae* bacteria, whereas after incubation with LPS of *E. coli* and heat-inactivated *S. aureus*, intact virus particles were clearly visible, confirming our previous titration data (Fig. 2A and 4A). A comparable direct effect of LPS on the virus particle has been determined before for influenza virus by TEM (15). Upon incubation with LPS for 20 min, it became difficult to detect the virus particles, since their structure had dramatically changed. Therefore, it is possible that the structures we considered as viruses, were in fact LPS- or bacteria-related structures rather than disrupted viruses.

In general, we determined an inhibitory trend of specific bacterial cell wall components on arbovirus infectivity *in vitro*. We now focused on the possible interactions between arboviruses and bacteria at the skin inoculation site. Another important factor is the mosquito saliva, which is also inoculated into the skin during the mosquito bite. It has been described that mosquito saliva can enhance arbovirus infection, dissemination and transmission with a negative effect on disease and mortality (42). Since we found that certain bacteria/ bacterial cell wall components can have a protective effect on arbovirus infectivity, it is thus possible that these two factors counteract and balance each other. It would therefore be of interest to study this in *in vivo* models in the future.

Another factor that might affect the interpolation of our *in vitro* results to patients, is the known individual variation in the composition of skin microbiota (5). In addition, it has been demonstrated that certain skin bacteria produce volatile organic compounds that attract or repulse *Aedes* mosquitoes (43). *Corynebacterium tuberculostearicum* and *Staphylococcus hominis* had a strong repellent effect and one of the bacteria that we tested, *C. amycolatum*, had a weak repellent effect (44). Since we determined that certain skin bacteria/bacterial cell wall components can have a protective anti-arboviral effect, this can imply that humans possessing higher abundancies of bacteria that can inhibit arboviruses or bacteria that can repel mosquitoes, could be more resistant to infections by certain arboviruses. On the other hand, it has been shown that dermal bacteria (which are most relevant for this research as arboviruses are inoculated into the dermis) are less affected by environmental factors, resulting in less variability among individuals (29) and thus making our results more generally applicable.

Together, our *in vitro* results are in line with recently published *in vivo* data (21, 22), which showed an aggravated arbovirus infection upon depletion of the host microbiota by administering antibiotics. These mouse studies confirmed that oral antibiotic treatment reduced bacterial colonization and altered bacterial populations in the faeces. Furthermore, antibiotics reduced host immune responses. A decrease in CD8+ T cells was measured in mice treated with antibiotics and infected with flaviviruses (WNV, DENV, ZIKV) (21). In addition, a decrease of type I IFN production and monocyte interferon stimulating gene expression due to the perturbation of the intestinal microbiota by antibiotic treatment resulted in enhanced alphavirus (CHIKV) infection (22). As mice were orally treated in these studies, the antibiotics affected the gut microbiota and caused a systemic effect. Until now, nothing was known about interactions between arboviruses and the host skin microbiota at the inoculation site. Our *in vitro* data on the direct interaction between host (skin) microbiota and viruses are a first step towards elucidation of this knowledge gap.

## Acknowledgments

We thank Elke Maas for excellent technical assistance. We thank Prof. K. Lagrou (UZ Leuven) for providing bacterial isolates and Prof. C. Drosten (Charité – Universitätsmedizin Berlin) for providing the CHIKV 899 strain. We thank the European virus archive, EVAg, supported by the European Union’s Horizon 2020 research and innovation programme under grant agreement No 871029, for providing MAYV TC625 and ZIKV SL1602, Suriname. We thank Dr. Els Vanstreels and Dr. Maarten Jacquemyn for their help with the set-up of the confocal fluorescence microscope.

This research was funded by a ZAP starting grant to LD by KU Leuven (STG/19/008). SJ is a fellow of the “Fund for Scientific Research” Flanders (FWO).

## Notes

### Competing Interest Statement

The authors have declared no competing interest.

